# Investigation of the Synergistic Effect of Enzymatic and Ultrasound-Induced Amyloid Microclot Degradation

**DOI:** 10.1101/2025.09.12.675963

**Authors:** Reza Rasouli, Brad Hartl, Soren Konecky

## Abstract

Amyloid microclots have been implicated in thrombotic complications across various pathological conditions such as Long COVID symptoms, yet their resistance to enzymatic fibrinolysis causes a therapeutic challenge. In this study we examine the effects of three fibrinolytic enzymes rtPA, Lumbrokinase, and Nattokinase on plasma-derived amyloid microclots, in combination with ultrasound-induced microstreaming and microbubbles. A lab-on-chip platform was used to expose the clots to ultrasound at 150, 300, and 500 kHz. Quantitative analysis revealed that ultrasound alone significantly disrupted clot structures, particularly at 150 kHz, where mean clot diameter was reduced by over 60% and large-clot count (>30 µm) dropped by more than 80% compared to controls. The addition of fibrinolytic enzymes, however, did not produce statistically significant effects at 150 or 300 kHz which indicates that mechanical forces were the dominant contributors to clot disruption. At 500 kHz, where ultrasound alone was less effective, enzymatic treatment moderately enhanced the reduction in large-clot burden. These results show the potential of low-frequency ultrasound as a primary method of amyloid microclot breakdown, with enzyme co-treatment offering limited but measurable effect.

## Introduction

The persistence of microvascular symptoms in post-infectious syndromes such as Long COVID has increased the interest in the role of amyloid-like fibrin structures known as fibrinaloid microclots. These aggregates are characterized by β-sheet–rich fibrin(ogen) conformations and are reported to be resistant to conventional enzymatic fibrinolysis^1–3^. Their presence has been documented in Long COVID via fluorescent dyes such as Thioflavin T (ThT), which selectively binds to misfolded fibrin and enables microscopic or flow cytometric visualization of the microclot burden ^4,5^. Similar microclot pathology has also been noted in Type 2 diabetes, myalgic encephalomyelitis/chronic fatigue syndrome (ME/CFS), and acute COVID-19, suggesting that amyloid fibrin aggregates may trigger broader chronic inflammatory or thrombo-inflammatory syndromes ^6–8^.

Microclots in Long COVID act as sinks for inflammatory and antifibrinolytic molecules. Proteomic analyses have revealed the entrapment of serum amyloid A (SAA), α2-antiplasmin, and clotting factors within these aggregates, forming a biochemically stable and pathologically active clot structure ^9,10^. Elevated levels of α2-antiplasmin inhibits activity plasmin which is the primary fibrinolytic enzyme and therefore prevent the breakdown of fibrin^11^. Consequently, conventional thrombolytic therapy such as rtPA often proves insufficient in treating amyloid clots, necessitating the search for alternative strategies.

Among the therapeutic candidates under investigation are two natural fibrinolytic enzymes called nattokinase and lumbrokinase. Nattokinase is a serine protease derived from *Bacillus subtilis* during the fermentation of soybeans (natto) and has demonstrated fibrinolytic and antithrombotic effects in vitro and in vivo ^12^. Lumbrokinase is a complex of proteolytic enzymes extracted from the earthworm *Eisenia fetida*, and exhibits more potent fibrinolytic activity, potentially acting through both plasminogen-dependent and independent pathways. These agents are being explored as alternatives or adjuncts to rtPA in managing fibrin-rich thrombi particularly due to their safety profiles and natural origin.

An additional strategy to overcome clot resistance involves the use of low-frequency ultrasound. Acoustic energy can enhance thrombolysis via mechanical mechanisms such as acoustic streaming, radiation force, and cavitation, which facilitate enzyme penetration and mechanically disrupt fibrin networks. ^13^ Studies combining ultrasound with microbubble contrast agents have shown further potentiation of thrombolytic efficacy, attributed to microbubble oscillation and collapse enhancing localized shear and clot porosity ^14–17^. Our recent investigation using amyloid clots demonstrated that low-frequency ultrasound (150 kHz) alone could significantly fragment microclots. In that study, we attributed the clot disruption to and stable cavitation and acoustic microstreaming. In response to acoustic pressure fields, microbubbles undergo volumetric oscillations which in turn generate high-velocity acoustic microstreams in the surrounding fluid. These microstreams impose localized shear forces on the clot structure and mechanically fragment amyloid microclots. This mechanical mechanism was sufficient to achieve substantial clot disruption under low-frequency ultrasound exposure.^18^

In this study we seek to evaluate the clot-lysis capacity of rtPA, nattokinase, and lumbrokinase in an *in vitro* model of amyloid microclots and their synergistic effect in the presence of low-frequency ultrasound and microbubbles. We assess whether enzymatic thrombolysis can be enhanced through acoustic exposure, and whether natural enzymes can match or exceed the efficacy of rtPA in lysing resistant clots. These findings aim to inform the development of novel therapeutic strategies for managing persistent fibrin pathology in Long COVID and related chronic thrombo-inflammatory conditions.

## Materials and Methods

### Plasma Preparation and Amyloid Microclot Formation

Porcine plasma was obtained from Innovative Research, Inc. (USA) and stored at -20 °C until use. To induce the formation of amyloid microclots, plasma aliquots (1 mL) were subjected to 12 consecutive freeze-thaw cycles, alternating between -20 °C and 37 °C for 2 hours per cycle.^18^ After completion of the cycles, samples were incubated at 37 °C to allow further amyloid aggregation and then diluted 1.5× prior to ultrasound treatment. This procedure was previously validated ^18^to generate Thioflavin T-positive amyloid microclots which mimick the resistant fibrinaloid aggregates observed in Long COVID and other chronic thrombo-inflammatory states.

### Fibrinolytic Enzyme Preparation

Three fibrinolytic agents were tested: recombinant tissue plasminogen activator (rtPA), lumbrokinase, and nattokinase. All enzyme solutions were prepared in phosphate-buffered saline (PBS) immediately before use. rtPA (Sigma-Aldrich, USA): Stock concentration was 0.1 mg/mL (300,000 IU/mL). A working solution of 3 μg/mL (∼900 IU/mL) was prepared by mixing 30 μL of rtPA stock with 970 μL of PBS. Lumbrokinase was obtained from Canada RNA Biochemical Inc., Richmond, Canada and reconstituted to ∼900 IU/mL. Nattokinase was purchased from GeneMill, Liverpool, UK and tested at 28 ng/ μL as per manufacturer suggestion. All enzyme preparations were gently mixed and kept on ice until addition to the LoC system.

### Microbubble Preparation

Vevo MicroMarker® contrast agents (FUJIFILM VisualSonics Corp., Japan) were used as exogenous microbubbles. Microbubbles were diluted in PBS according to the manufacturer’s protocol immediately prior to each experiment and added to the clot-containing solution prior to ultrasound exposure.

### Lab-on-a-Chip (LOC) Device Fabrication

The experimental platform was built using a custom-fabricated lab-on-a-chip (LOC) device designed to simulate a 6 mm diameter popliteal vein, chosen for its clinical relevance in transcutaneous ultrasound targeting. The molds were created using Onshape CAD software (PTC Inc., USA) and manufactured via digital light processing (DLP) 3D printing. Channels were cast in Momentive RTV615 silicone elastomer and bonded to a 20 μm silicone sheet (Gteek, UK) using plasma surface activation.

### Ultrasound Exposure Setup

Experiments were conducted under static conditions inside a custom-designed acoustic water bath, housing the LOC device, piezoelectric transducers, and mounting components. Transducers were placed at the base of the bath, and the LOC was held at the pre-measured acoustic focal plane to ensure consistent energy delivery. Degassed water served as the acoustic coupling medium. Ultrasound was applied at three frequencies: 150 kHz, 300 kHz, and 500 kHz, in continuous wave mode. A function generator (DG4162, Rigol Technologies, China) supplied the waveform, and an amplifier (1040L, Electronics & Innovation, USA) controlled voltage delivery. The mechanical index (MI) was maintained at 0.3 across all frequencies by adjusting acoustic pressure (PNP).

### Fluorescent Labeling and Imaging of Microclots

To confirm amyloid fibrin structure, Thioflavin T (ThT) staining was performed. A 0.03 mM ThT solution was prepared in PBS (pH 7.4). Samples were incubated at room temperature for 30 minutes after treatment in the dark prior to imaging. Imaging was performed using fluorescence microscopy with appropriate excitation/emission filters (450–530 nm for ThT). Images were analyzed to quantify clot presence, size, and fragmentation post-treatment.

### Clot Lysis Assessment and Image Analysis

Clot breakdown was quantified by microscopic image analysis, comparing microclot size distributions and particle counts before and after treatment. Particular emphasis was placed on the reduction in large (>30 μm) amyloid microclots, which have been associated with pathological vascular occlusion in Long COVID. Image J was used to calculate mean clot size and total clot area per field of view.

### Statistical Analysis

Statistical significance was assessed using Brown-Forsythe/Welch’s ANOVA followed by Dunnett’s T3 post hoc test for multiple comparisons. Significance thresholds were defined as: *p* < 0.05 (\**), p < 0*.*01 (*\*\**), p < 0*.*001 (*\*\*\*), and *p* < 0.0001 (****).

## Results

### Effect of rtPA, Ultrasound, and Microbubbles on Amyloid Microclot Diameter

To evaluate the efficacy of the enzymes and in combination with ultrasound and microbubbles on the breakdown of amyloid microclots, we measured the average clot diameter and number of large microclots (>30 µm) under various experimental conditions.

### Effect of Ultrasound on Microclot lysis

Figure 1A shows fluorescence microscopy images of green (ThT)-stained amyloid microclots exposed to ultrasound at varying frequencies in the presence of microbubbles. The images show fragmentation of microclots under ultrasound. At 150 kHz, the microclots appeared highly dispersed, with smaller, less fluorescent aggregates visible compared to that of the control, indicating substantial mechanical disruption. In contrast, while clot dispersion was still evident at 300 kHz and 500 kHz, more intact larger clots remained visible which agrees with our previous report.^18^ These findings are quantified in Figure 4B which reports the mean clot diameters and number of large clots across the four conditions. In untreated control samples, mean microclot diameters is ∼17.5 µm, with minimal variance across replicates. Application of ultrasound in the presence of microbubbles significantly reduced mean diameters in a frequency-dependent manner. At 150 kHz, average diameters dropped to ∼8 µm. At 300 kHz and 500 kHz, clot sizes were moderately reduced (∼10–11 µm), confirming the mechanical lysis. Figure 4B (right) shows the number of large microclots (>30 µm) in each condition. The untreated control exhibited a high burden of large aggregates (mean count ≈ 275), while exposure to 150 kHz resulted in a >80% reduction to ∼50 microclots. Both 300 kHz and 500 kHz also significantly reduced the number of large clots although the reduction was less pronounced compared to 150 kHz.

**Figure 1.**
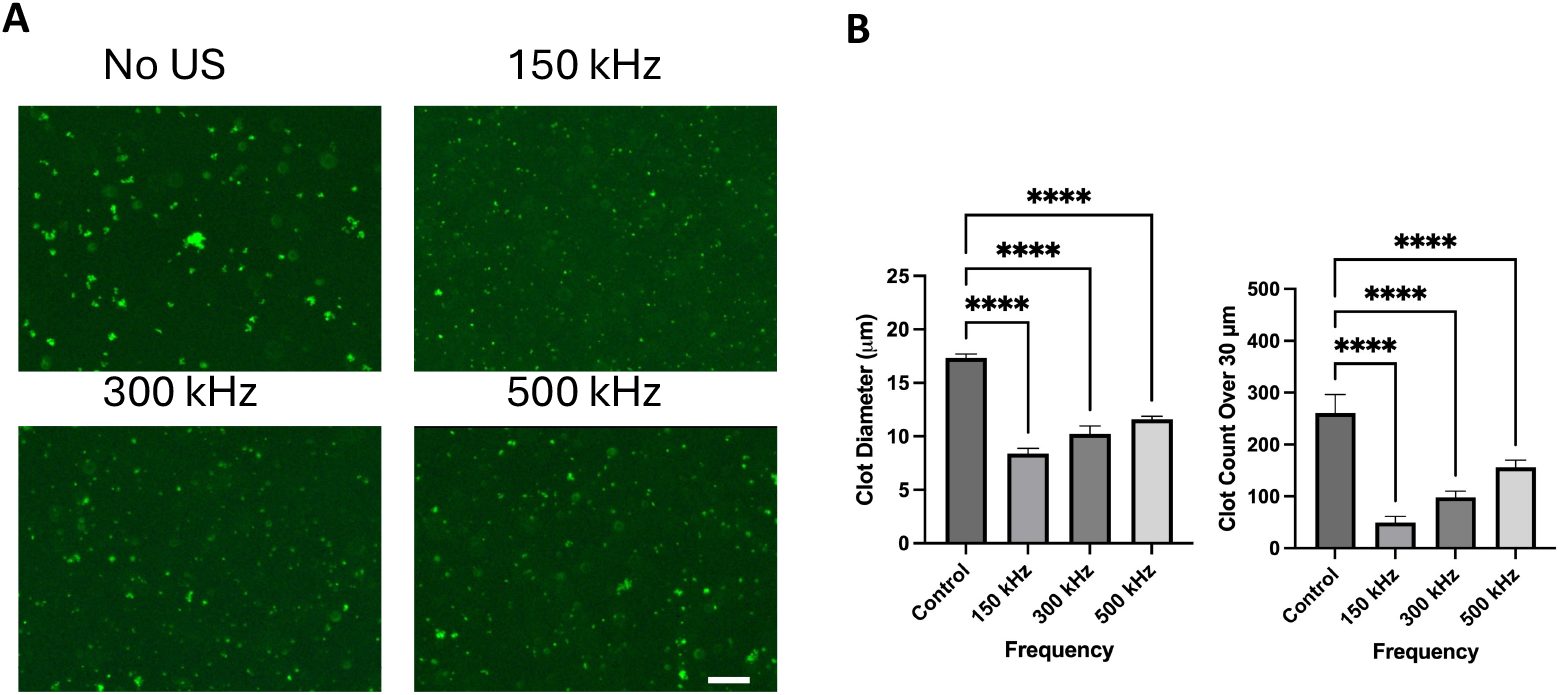
Effect of ultrasound on clot lysis. A) Thioflavin T-labeled microclots after control (No US) and US exposure at 150, 300, and 500 kHz (scale bar = 50 µm). B) quantification of clot diameter (µm) (left) and number of large clots (right); error bars indicate SD.

### Effect of rtPA on Microclot lysis

To observe the enzymatic effect of on clot lysis, we repeated the experiment with the addition of rtPA at a concentration of 900 IU/mL. Figure 2A shows fluorescence microscopy images of ThT-stained microclots treated with rtPA and microbubbles across a range of ultrasound frequencies. In the rtPA only condition (no US), abundant intact microclots remained visible, indicating that rtPA alone had minimal fibrinolytic activity under static conditions. Upon exposure to 150 kHz ultrasound, a significant reduction in clot size was observed. At 300 kHz and 500 kHz, clot breakdown remained evident, but larger aggregates persisted more frequently, suggesting that the synergistic effects of ultrasound and rtPA were strongest at lower frequencies.

**Figure 2.**
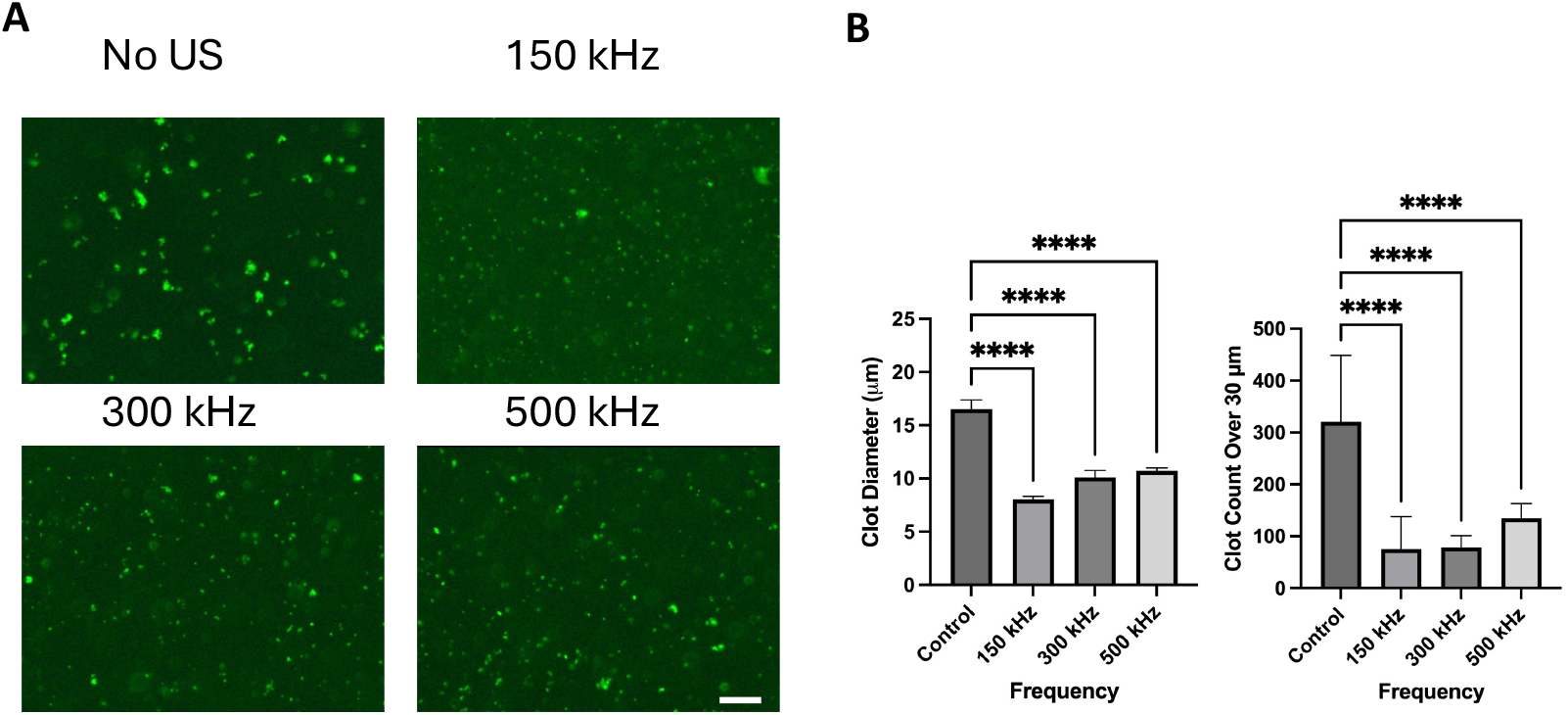
Effect of rtPA and ultrasound on clot lysis. A) shows fluorescent image of ThT-labeled microclots after rtPA only control and rtPA plus US exposure at 150, 300, and 500 kHz (scale bar = 50 µm). B) quantification of clot diameter (left) and number of large clots (right). Error bars indicate SD.

Quantitative results are shown in Figure 2B. In the left panel, mean clot diameter under control conditions remained elevated (∼17 µm). Application of 150 kHz ultrasound significantly reduced the average clot size to ∼8 µm. At 300 kHz and 500 kHz, diameters were moderately reduced (∼10 µm), further confirming the frequency-dependent potentiation of enzymatic fibrinolysis. The number of large clots dropped sharply from (∼ 300 in the control group to fewer than 100 following 150 kHz treatment which shows a substantial decrease in large-clot burden.

### Effect of Lumbrokinase on Microclot lysis

We further tested lumbrokinase (900 IU/mL) to evaluate its ability to degrade amyloid microclots when combined with ultrasound and microbubbles. Figure 3A shows that in the absence of ultrasound (Lumbrokinase only), microclots remained intact, indicating minimal lytic activity by lumbrokinase under static conditions. Upon application of 150 kHz ultrasound, a clear reduction in clot size and density was observed, with fewer and smaller fluorescent structures. At 300 kHz and 500 kHz, clot disruption was still apparent, but the presence of larger microclots suggests reduced efficacy at higher frequencies.

**Figure 3.**
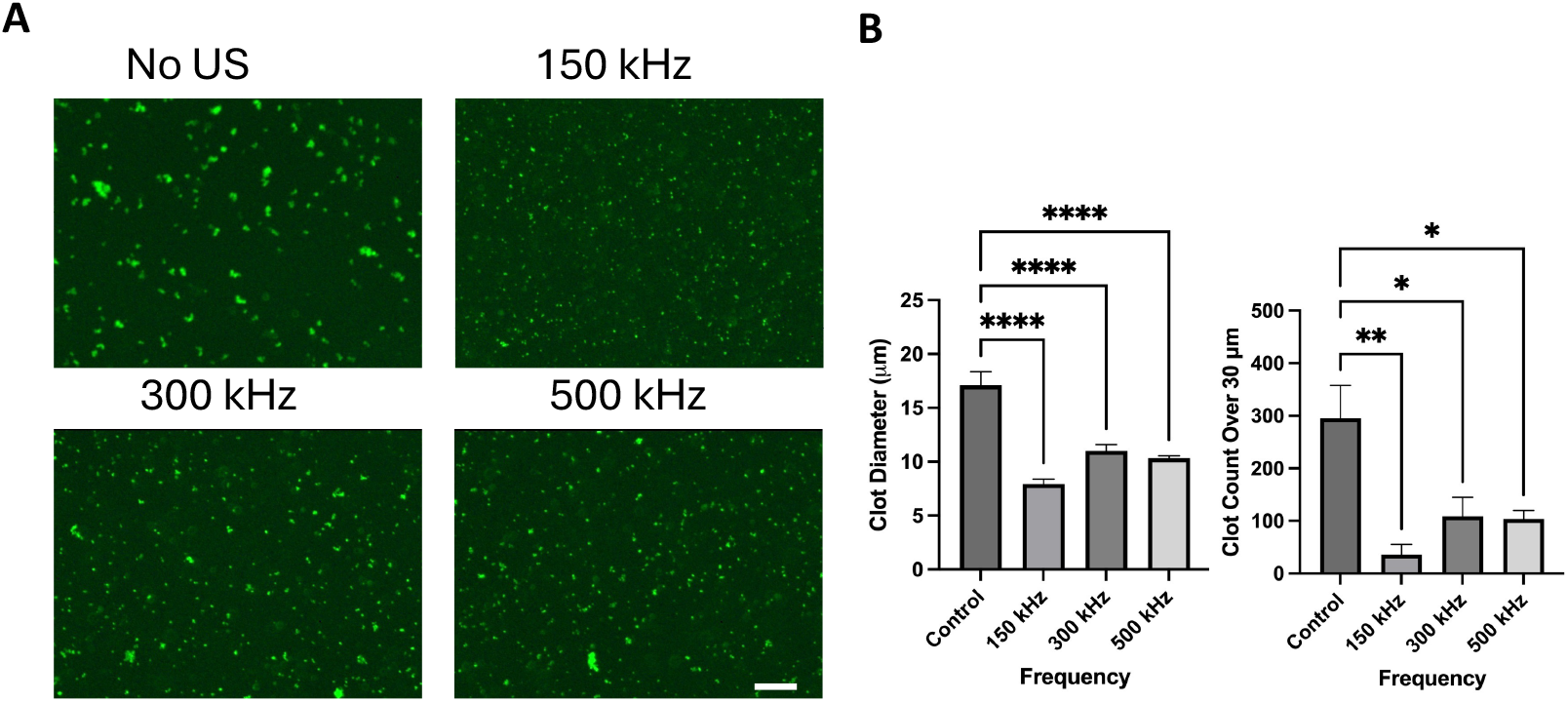
Effect of lumbrokinase and ultrasound on clot lysis. A) shows fluorescent image of ThT-labeled microclots after lumbrokinase only control and lumbrokinase plus US exposure at 150, 300, and 500 kHz (scale bar = 50 µm). B) quantification of clot diameter (left) and number of large clots (right). Error bars indicate SD.

**Figure 4.**
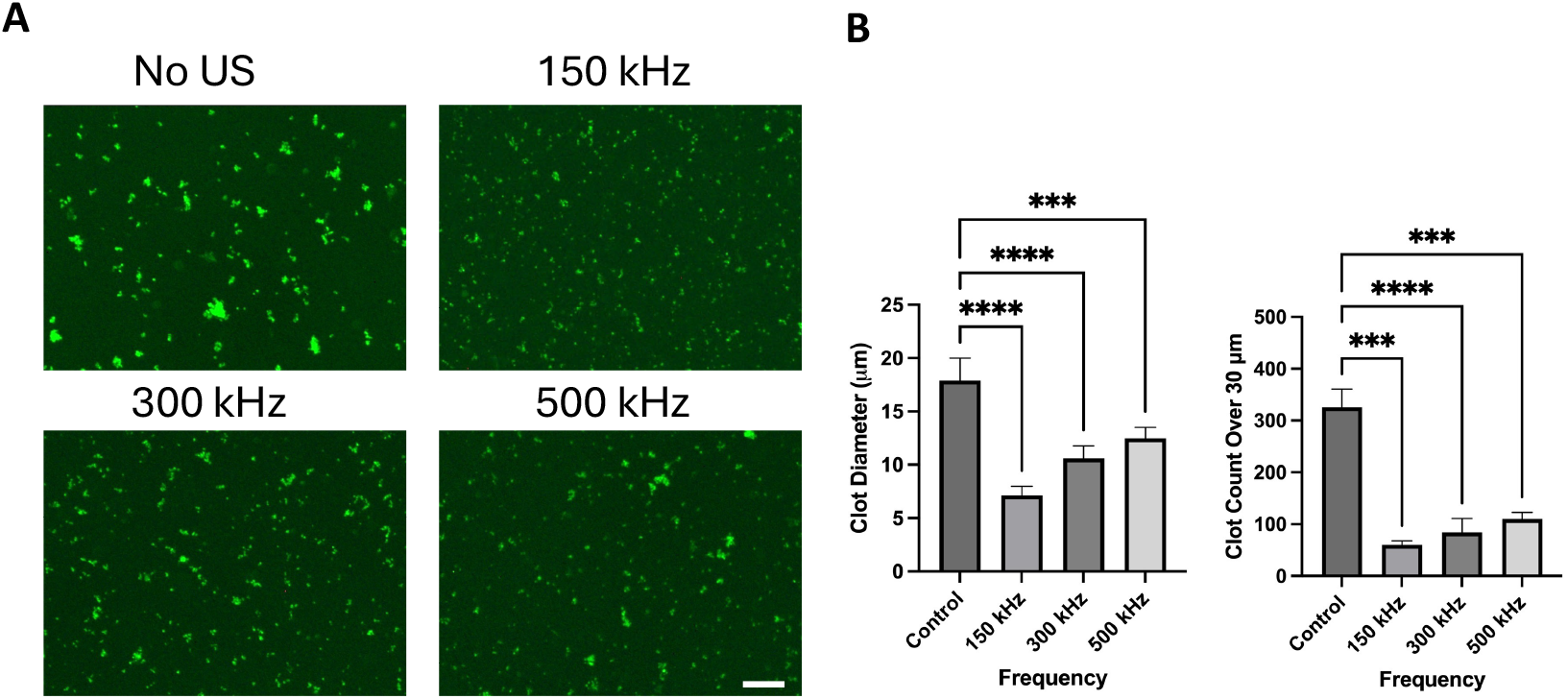
Effect of nattokinase and ultrasound on clot lysis. A) shows fluorescent image of ThT-labeled microclots after nattokinase only control and nattokinase plus US exposure at 150, 300, and 500 kHz (scale bar = 50 µm). B) quantification of clot diameter and number of large clots (right). Error bars indicate SD.

**Figure 5.**
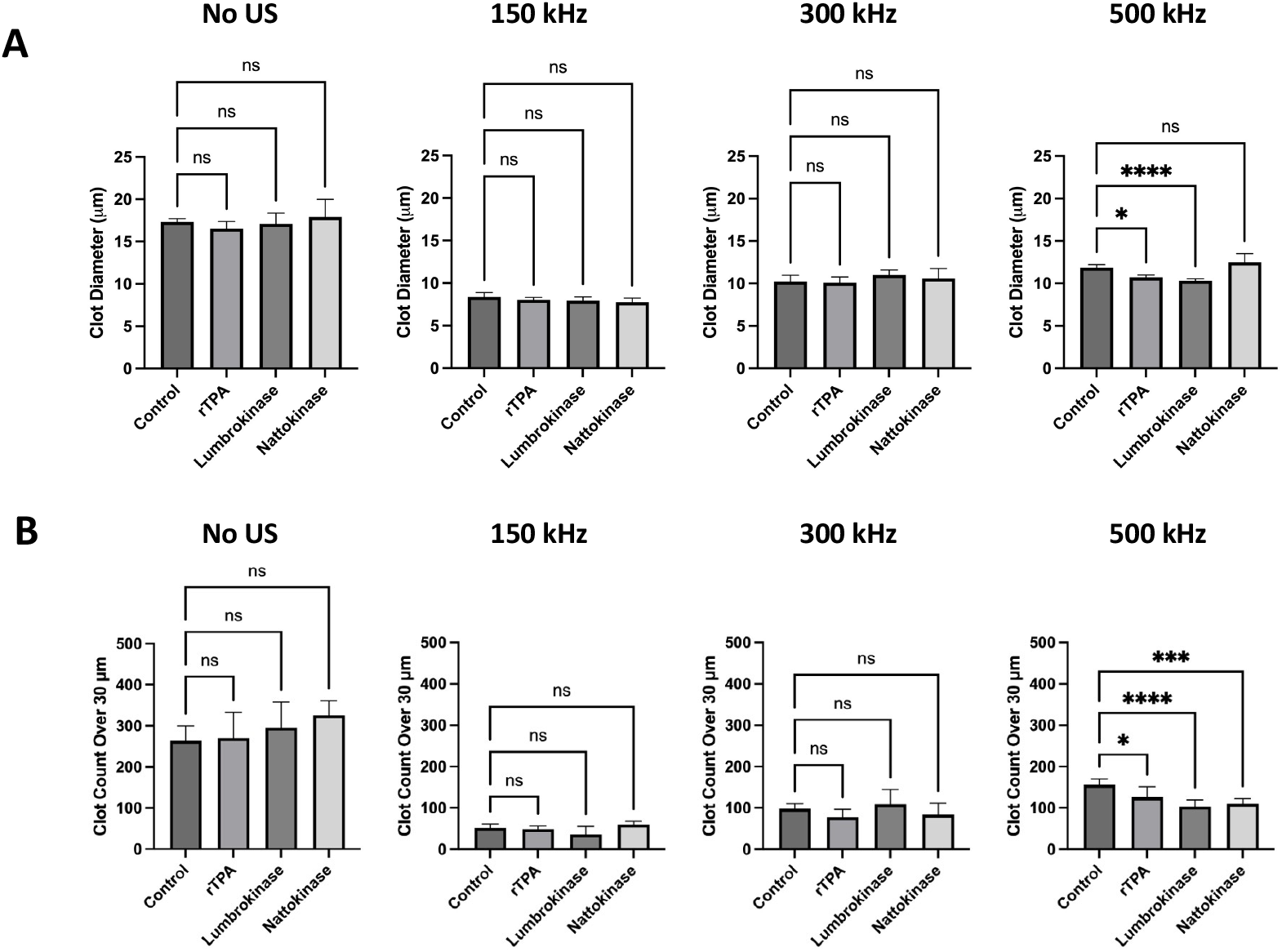
Comparison of clot lysis between enzymes at different ultrasound (US) frequencies. Control samples received no US exposure, while treated samples were exposed for 1 min at the indicated frequencies. Error bars represent SD.

Figure 3B shows that the mean clot diameter in control samples was approximately 17 µm, which decreased significantly to ∼7–8 µm at 150 kHz. At 300 kHz and 500 kHz, average diameters remained moderately low (∼10–11 µm). The large clot quantification followed previous results in that a significant dropped substantially at 150 kHz was observed while showing less effect at 300 kHz and 500 kHz.

### Effect of Nattokinase on Microclot lysis

Finally, we tested nattokinase to assess its clot lysis potential in combination with ultrasound and microbubbles. Similar pattern as other enzymes were observed where the Nattokinase alone did not substantially improve clot lysis in the absence of ultrasound, as average diameters remained high (∼18 µm) while at 150 kHz, clot diameter were visibly reduced. Although clot breakdown was still observable at 300 kHz and 500 kHz, a greater number of residual large clots persisted compared to 150 kHz.

Quantitative measurements show that mean clot diameter in untreated controls was ∼18–19 µm, decreasing significantly to ∼8 µm at 150 kHz. At 300 kHz and 500 kHz, diameters remained modestly reduced (∼11–12 µm), suggesting reduced ultrasound efficacy at higher frequencies. The right panel shows the number of clots exceeding 30 µm, which dropped sharply from controls to fewer than 100 at 150 kHz. Frequencies of 300 kHz and 500 kHz also reduced large-clot burden, but to a less extent.

### Comparison to Controls

To evaluate the relative performance of each enzyme across ultrasound conditions, we compared microclot diameters under no ultrasound (static) and at 150 kHz, 300 kHz, and 500 kHz, with each frequency grouping including a control and all three enzyme treatments.

Under no ultrasound condition, none of the enzymes significantly reduced clot size relative to the control. Mean clot diameters remained in the range of 17–18 µm across rtPA, nattokinase and lumbrokinase. This confirms that without acoustic stimulation, enzymatic degradation of amyloid microclots is minimal, consistent with their known fibrinolytic resistance and dense amyloid content. Likewise, under 150 kHz and 300 kHz sonication, the addition of rtPA, lumbrokinase, and nattokinase did not have a statistically significant effect on the mean microclot diameter. At 500 kHz, treatment with either rtPA or lumbrokinase resulted in a significant decrease in clot diameter compared to control, whereas the effect of nattokinase was not significant.

The number of clots larger than 30 µm also showed similar trend and in the absence of ultrasound there were no significant differences between control and any enzymatic treatment which suggust that rtPA, lumbrokinase, and nattokinase had minimal activity on their own. Similarly, at 150 kHz and 300 kHz, clot counts remained statistically unchanged across groups. However, at 500 kHz, all enzymes showed a significant reduction in large-clot count compared to the control. rtPA achieved a modest effect, while lumbrokinase and nattokinase showed a more significant reduction.

## Discussion

The findings show the predominant role of ultrasound mechanical forces in amyloid microclot disruption. Among the tested conditions, 150 kHz ultrasound consistently produced the most substantial reduction in both microclot diameter and large-clot number, independent of the enzymatic treatment applied. These observations are consistent with our previous study, where ultrasound-induced clot lysis occurred in the absence of enzymatic agents, and mechanical forces from acoustic microstreaming were identified as key contributors to disruption. Microbubble oscillations generated intense local shear forces at the clot interface which led to fragmentation of otherwise resilient amyloid microclots. ^18^

Across all enzymes tested there was limited clot lysis in the absence of ultrasound. Furthermore, enzymatic enhancement at lower frequencies (150 and 300 kHz) was not statistically significant. However, at 500 kHz, where mechanical disruption was less pronounced compared to lower frequencies, we observed a modest but consistent improvement in clot reduction with the addition of enzymes. This suggests when mechanical disruption is less dominant, enzymatic activity becomes more detectable. One plausible explanation is that acoustic streaming at this frequency enhances convective transport of enzymes into the interior of partially disrupted clots, increasing access to cleavage sites that are otherwise sterically shielded. Conversely, at lower frequencies ultrasound microstreams and their shear forces dominate, clots may be fragmented to the limit that additional enzymatic degradation either contributes little to further size reduction or produces soluble fragments below the detection limit of current microscopy. In such cases, the contribution of enzymes may be real but not measurable, as degraded products disperse beyond detection thresholds. Nonetheless, the findings suggest that the mechanical effects induced by low-frequency ultrasound are effective enough to overshadow additional enzymatic contribution in clot degradation. Enzymatic treatments alone in the absence of ultrasound failed to produce significant clot lysis which suggest the essential role of mechanical degradation of amyloid-rich microclots.

One explanation for the low performance of enzymes, particularly rtPA, compared to conventional clot studies may lie in the fundamentally different composition of the freeze–thawed microclots, as opposed to the more commonly employed fibrin clots. Standard fibrin networks polymerized by thrombin are relatively porous and readily cleaved by plasmin after rtPA activation, whereas amyloid-rich microclots are abnormally dense aggregates enriched in cross-β sheet structures which can block plasmin cleavage sites and reduce the efficacy of rtPA-mediated fibrinolysis.^19^ This phenomenon aligns with prior reports that amyloid microclots in COVID patients show resistance to enzymatic lysis.^20^ Proteomic analyses of such clots have further revealed the enrichment of antifibrinolytic proteins including α2-antiplasmin, platelet factor 4, and serum amyloid A, which likely suppress plasmin generation and activity.^10,20^

## Conclusion

This study demonstrates that low-frequency ultrasound with microbubbles, particularly at 150 kHz and 300 kHz, effectively reduces amyloid microclot size and burden through mechanical mechanisms, independent of enzymatic degradation. In contrast, fibrinolytic enzymes including rtPA, lumbrokinase, and nattokinase showed minimal clot-dissolving activity under no US conditions and their contribution to clot lysis was statistically insignificant when combined with ultrasound at lower frequencies. However, at 500 kHz, where mechanical disruption was less pronounced, enzymatic treatments produced a measurable reduction in large-clot burden. These findings suggest that the efficacy of enzymatic fibrinolysis is dependent on the acoustic environment, and that mechanical forces are the predominant factor in microclot disruption under the tested conditions. The integration of ultrasound and fibrinolytic agents may represent a viable approach for targeting enzyme resistant amyloid microclots in thromboinflammatory disorders such as Long COVID.

